# Genomic architecture of the major histocompatibility complex genes in the Neotropical Midas cichlid

**DOI:** 10.64898/2026.11.04.764204

**Authors:** Juan S. Echeverry-Pérez, Marta Barluenga

## Abstract

The Major Histocompatibility Complex (MHC) plays a central role in vertebrate adaptive immunity, yet its genomic architecture remains poorly characterized in many evolutionary model systems, including the Neotropical Midas cichlid *Amphilophus citrinellus*. Here, we integrate chromosome-level functional genome annotation, transposable element (TE) profiling, transcriptomics, phylogenetics, molecular selection analyses, and protein structural modeling to characterize its MHC repertoire. We identified 298 MHC-related loci across 18 chromosomes, including 30 MHC-I genes, 15 MHC-IIA genes, and 20 MHC-IIB genes, together with antigen-processing and regulatory components. MHC-I and MHC-II genes were located on different chromosomes, consistent with the MHC non-linkage pattern characteristic of many teleosts. Gene duplication and phylogenetic patterns support birth-and-death evolution, resulting in a structurally diverse repertoire with broadly conserved protein domains. TE distribution varied across MHC genomic regions, suggesting a potential contribution to genomic diversification. Molecular selection analyses detected positive selection in MHC-I and IIB genes, whereas MHC-IIA showed more limited evidence of positive selection. Expression patterns varied across tissues, with relatively high expression of several MHC genes in the spleen and thymus, while structural models retained characteristic MHC folds. Phylogenetic analyses placed previously characterized MHC sequences with broader multilocus repertoire, suggesting that population level diversity may reflect both allelic polymorphism and variation among paralogous loci. Together, these findings provide a comprehensive chromosome-resolved characterization of the MHC repertoire in a Neotropical cichlid and establish a genomic framework for investigating immune gene diversification during the Midas cichlid radiation.

**Significance Statement:** The MHC is central to vertebrate adaptive immunity and among the most evolutionarily dynamic gene families, yet its genomic architecture remains unresolved in many non-model vertebrates. We show that MHC diversity in the Midas cichlid extends beyond the allelic variation previously characterized, encompassing an extensive, chromosomally structured repertoire of duplicated loci. By integrating genome annotation, transposable element profiling, transcriptomics, structural modeling, phylogenomics, and molecular selection analyses we reveal patterns of gene duplication, evolutionary diversification, and structural conservation that have shaped the MHC repertoire. This work shifts MHC studies in Midas cichlids from a sequence-based to a genome-wide perspective and provides a foundation for interpreting MHC diversity in natural populations and investigating the evolution of adaptive immunity in teleost fishes.

## Introduction

The major histocompatibility complex (MHC) is a cornerstone of the adaptive immune system in jawed vertebrates, where it plays a central role in antigen presentation to T cells and links pathogen recognition to immune effector responses (Germain 1994; Zinkernagel and Doherty 1997; Pinesha et al. 2022). MHC class I (MHC-I) and class II (MHC-II) molecules primarily present peptides derived from intracellular and extracellular antigens, respectively (Janeway et al. 2001; Murphy and Weaver 2016). Antigen presentation also relies on a network of associated genes involved in peptide processing and transport, MHC assembly, and transcriptional regulation, including *b2m*, *tap*, *tapbp*, *psmb*, *cd74*, *ciita* and *rfx* family members (Krawczyk et al. 2004; Seguín-Estévez et al.2009; Pinesha et al. 2022).

Beyond their genetic diversity, MHC proteins exhibit remarkable conformational plasticity, enabling them to bind a wide range of antigenic peptides with high specificity (Wieczorek et al. 2017). Because of their central role in pathogen recognition, MHC genes often evolve more rapidly than most genomic regions, allowing hosts to maintain a dynamic equilibrium with changing parasite and microbial communities (Neefjes et al. 2011; Eizaguirre et al. 2012). This accelerated evolution is characterized by extreme polymorphism and frequent gene duplication, making the MHC one of the most complex gene families in vertebrate genomes (Geraghty et al. 2002). Consequently, these loci have often been regarded as “speciation genes” because of their potential role in reproductive isolation and adaptive diversification (Eizaguirre et al. 2009; Malmstrøm et al. 2016). Together, MHC and MHC-associated genes form an integrated antigen–presentation system whose genomic organization and diversification remain incompletely understood outside a few model vertebrates.

In mammals, the MHC-I and MHC-II genes are typically organized within a single predominant, tightly linked genomic region (Shiina et al. 2017), with additional paralogous regions containing duplicated genes (Kasahara 1999). In teleost fishes, MHC-I and MHC-II loci are generally located on different chromosomes, a condition known as MHC non-linkage (Sato et al. 2000; Michalová et al. 2000). This reorganization likely reflects the combined effects of the two early vertebrate whole-genome duplications (Sidow 1996; Meyer and Schartl 1999) and the teleost-specific third round of duplication (Meyer and Van de Peer 2005). Subsequent clade-specific rearrangements, losses, and local expansions, have further shaped MHC architecture, resulting in pronounced copy-number variation, multilocus organization, and lineage diversification (Yamaguchi and Dijkstra 2019). In teleosts, MHC-I genes are currently classified into six major lineages: U, Z, S, L, P, and H (Dijkstra et al. 2007; Grimholt et al. 2015, 2019). MHC-II genes have been classified into three major lineages DA, DB, and DE, based on sequence characteristics and phylogenetic relationships (Dijkstra et al. 2013;; Grimholt and Lukacs 2021).

Cichlid fishes provide an especially powerful system for examining these evolutionary dynamics. They are iconic models for studying adaptive radiation, explosive speciation, ecological diversification, and remarkable genomic plasticity (Barluenga et al. 2006; Salzburger 2018; Kautt et al. 2020; Ronco et al. 2021). The MHC has emerged as a promising framework for understanding how immunity may contribute to adaptation, local divergence, and potentially reproductive isolation (Blais et al. 2007; Eizaguirre et al. 2009, 2012; Hablützel et al. 2016). Over the past decades, MHC characterization in cichlids has primarily focused on MHC-IIB loci, revealing both conserved and clade-specific features across various geographic regions (Ono et al. 1993; Málaga-Trillo et al. 1998; Figueroa et al. 2000; Hablützel et al. 2014). This focus on MHC-IIB has been largely driven by the high polymorphism and functional relevance of exon 2, which encodes the peptide-binding β1 domain, together with the feasibility of targeting this region using PCR-based approaches in population-level studies. However, such targeted approaches provide limited information on the number, identity and genomic location of the MHC loci being amplified. In Neotropical cichlids, considerable effort has been directed toward the Midas cichlid species complex (*Amphilophus* spp.), where previous studies revealed high allelic diversity and signals of balancing and non-parallel selection, but could not distinguish allelic polymorphism from locus duplication or assess the full genomic architecture of the system (Hofmann et al. 2017; Bracamonte et al. 2022; Lozano-Martín et al. 2023).

To address these knowledge gaps, we used the high-quality, haplotig-purged chromosome-level reference genome of the Neotropical Midas cichlid (*Amphilophus citrinellus*) to characterize the MHC genomic architecture and organization. Specifically, we (i) characterized the complete MHC repertoire, including MHC-I, MHC-IIA, and MHC-IIB genes, as well as antigen-processing components, transcriptional regulators, and associated genes, by identifying gene loci, copy numbers, and intron–exon organization; (ii) assessed gene structural integrity, protein domain architectures, and the distribution of transposable elements (TEs); (iii) assigned MHC-I and MHC-II genes to phylogenetic lineages and reconstructed phylogenetic relationships; (iv) characterized expression patterns across tissues; and (v) evaluated molecular selection pressures using codon-based d_N_/d_S_ models, and predicted three-dimensional structures of selected MHC proteins. Together, these analyses provide a comprehensive genomic and evolutionary characterization of the MHC repertoire, establishing a genomic framework for investigating immune gene diversification and its potential relevance to local adaptation and rapid evolutionary diversification in Neotropical cichlids.

## Results

### Overview and structural quality of the annotated Midas cichlid genome

We used the haplotig-purged, chromosome-level reference assembly of the Midas cichlid (*Amphilophus citrinellus;* GCA_013435755.1; Kautt et al. 2020) to characterize the MHC gene repertoire while minimizing the risk of interpreting allelic variants as duplicated loci. The assembly spans 0.96 Gb across 24 primary chromosomes (25.99–62.47 Mb), with a GC content of 40.96%, an N50 of 37.15 Mb, and an L50 of 12 (Table S1).

Structural and functional genome annotation, supported by transcriptomic data from multiple tissues and developmental stages (Table S2), identified 29,581 total genes, including 25,925 protein-coding genes, 2,721 non-coding genes, and 935 pseudogenes (Table S3). Gene-space completeness was high, as indicated by BUSCO (99.6%; Actinopterygii_odb10) and OMArk (95.04% consistency with Cichlidae HOGs; 98.17% species match). Proteome-wide quality assessment indicated high protein-coding probability for 99.1% of predicted models and intact reading frames in 99.92% of transcripts (Table S4).

Genome-wide transposable element (TE) annotation revealed that repetitive sequences accounted for 41.2% of the assembly, with DNA transposons (∼14.8%) and LINEs (∼11.1%) representing the largest fractions, followed by unclassified TEs (∼7.3%), LTR retrotransposons (∼4.3%), and other repetitive elements (∼2.7%), Rolling-circle elements (∼0.48%), SINEs (∼0.32%), and Penelope-like elements (∼0.2%) accounted for smaller proportions (Table S5).

### Genomic organization, repertoire composition, structural variation, and nomenclature of MHC genes

Combining functional genome annotation with homology searches against human MHC reference sequences (Supplementary file S1) and curated Neotropical and African cichlid sequences (Supplementary file S2), we identified 298 MHC-related gene copies distributed across 18 chromosomes in the Midas cichlid genome. These copies were assigned to distinct genomic loci based on their genomic coordinates and structural variation. The analysis included MHC-I and MHC-II genes, antigen processing components (*tap*, *psmb*, and *b2m* members), transcriptional regulators (*ciita* and *rfx* members), and additional associated genes (*brd*, *cd74*, and *mHA* members). The identified gene models were further evaluated using structural and sequence quality criteria (Table S6 and S7).

In humans, the MHC is primary organized on chromosome 6, comprising the Extended Class II, Class II, Class III, Class I, and Extended Class I regions (Shiina et al. 2017). In contrast, MHC-related genes in the Midas cichlid were distributed across 18 chromosomes (Fig. S1), with major concentrations on chromosomes 3, 4, 5, 8, 11, 17, and 22 (Fig. 1). These regions contained multiple MHC-I and MHC-II gene copies, particularly on chromosomes 3, 17 and 22, highlighting differences in gene copy number and genomic organization relative to the human MHC.

**Fig. 1.**
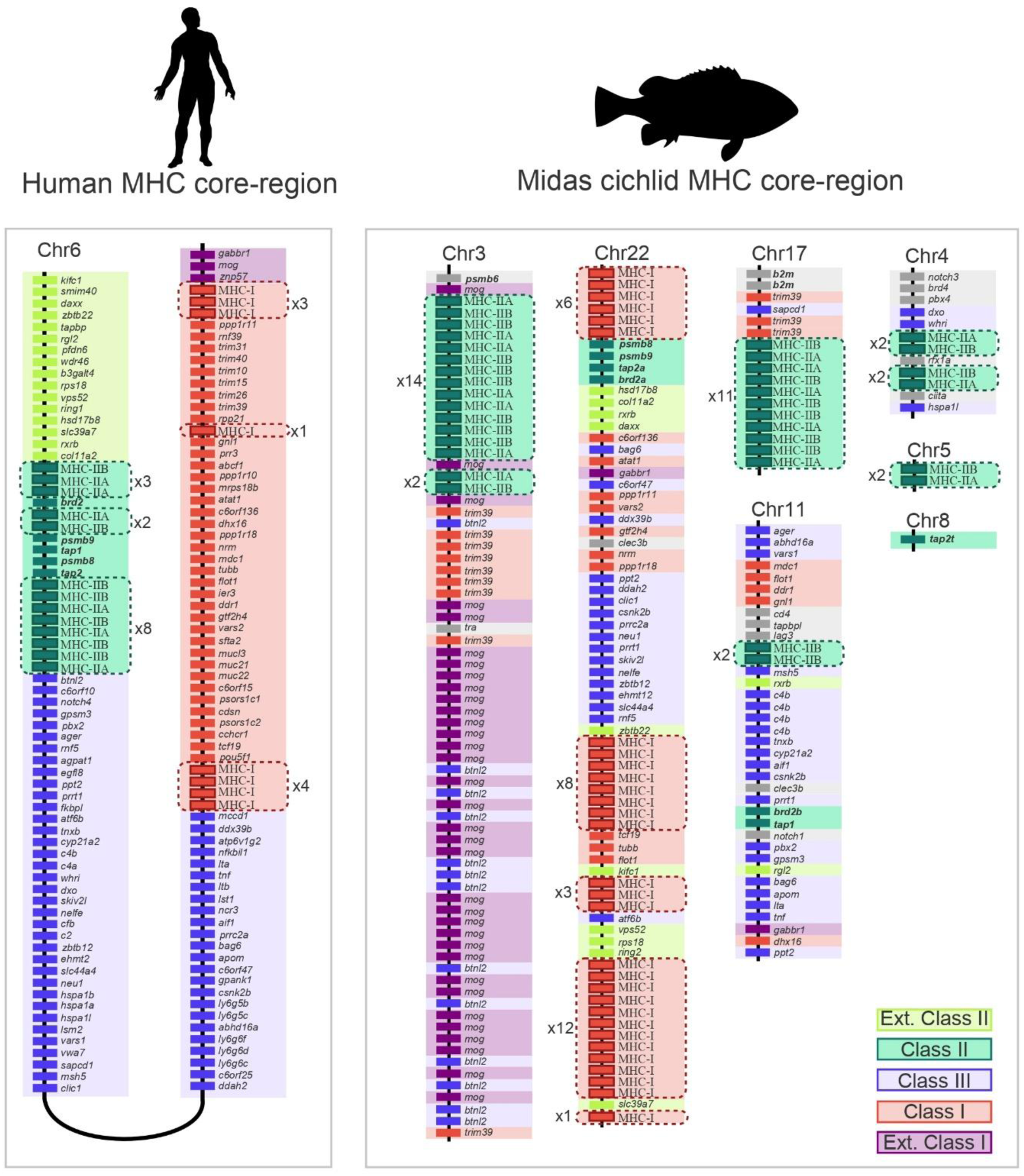
Genomic organization of MHC-related loci in humans and the Midas cichlid. Schematic comparison of the human MHC region on chromosome 6p21.3 and MHC-related regions across seven chromosomes in the Midas cichlid. Dashed boxes highlight MHC-I and MHC-II gene duplications, with xN indicating the number of gene copies. An expanded version is provided in Fig. S1.

Maximum likelihood (ML) phylogenetic analyses assigned MHC-I and MHC-II sequences to established vertebrate MHC lineages (Table S8 and S9; Supplementary files S3–S5). The MHC-I repertoire, located exclusively on chromosome 22, comprised 30 genes, 45 transcripts, and one pseudogene, alongside associated genes (*psmb8*, *psmb9*, *tap2a*, and *brd2a*) (Fig. 2). Most MHC-I sequences belonged to the U lineage (Amci-U1 to Amci-U27), while two were assigned to the Z lineage (Amci-Z1 and Amci-Z2) and one to the H lineage (Amci-H1). No sequences were assigned to the S, L, or P lineages (Fig. S2 and Table S9).

**Fig. 2.**
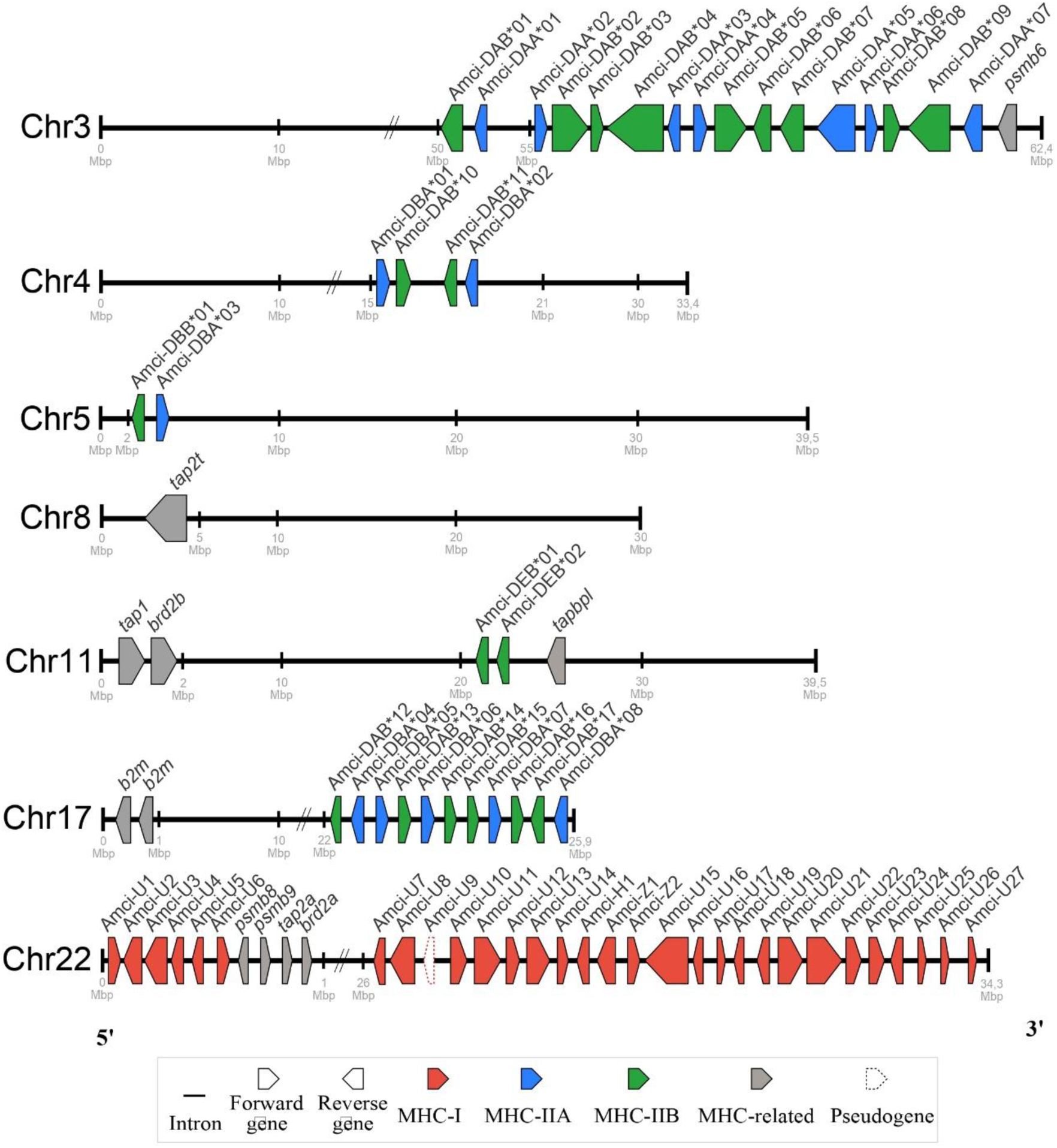
Genomic distribution of the MHC genes in the Midas cichlid. Schematic representation of MHC-I, MHC-IIA, MHC-IIB and associated genes across seven chromosomes. Arrows indicate gene position, orientation, and relative length, with colors distinguishing gene families and dashed outlines indicating pseudogenes. Chromosomal coordinates are expressed in megabases (Mb), and discontinuities indicate large omitted intervals.

MHC-II genes and associated components (*b2m*, *psmb6*, *tap1*, *tap2t*, *tapbpl*, and *brd2b*) were distributed across chromosomes 3, 4, 5, 8, 11, and 17 (Fig. 2). The MHC-IIA repertoire comprised 15 genes and 18 transcripts. Genes on chromosome 3 were assigned to the DA lineage (Amci-DAA*01 to Amci-DAA*07), whereas those on chromosomes 4, 5, and 17 belonged to the DB lineage (Amci-DBA*01 to Amci-DBA*08). No MHC-IIA sequences were assigned to the DE lineages (Fig. S3 and Table S9).

The MHC-IIB repertoire comprised 20 genes and 25 transcripts. Genes on chromosome 3 clustered within the DA lineage (Amci-DAB*01 to Amci-DAB*09), while those on chromosomes 4 and 17 formed a distinct DA subclade (Amci-DAB*10 to Amci-DAB*17). The gene on chromosome 5 was assigned to the DB lineage (Amci-DBB*01), and those on chromosome 11 to the DE lineage (Amci-DEB*01 and Amci-DEB*02) (Fig. S4 and Table S9).

MHC genes exhibited substantial structural variation in gene length, coding sequence (CDS) length and exon-intron organization (Fig. 3 and Table S10). MHC-I genes showed the broadest range of exon numbers (3–8), with gene lengths ranging from 3,969 to 52,428 bp. MHC-IIA genes were comparatively more uniform, spanning 1,752–14,617 bp with 4–6 exons, while MHC-IIB genes showed the widest range of gene lengths (1,065–64,445 bp) and contained 4–8 exons. Intron lengths generally showed greater relative variation than exon lengths, although substantial variation was also observed in some exons. Antigen-processing and regulatory genes generally showed less pronounced structural variation (Table S10).

**Fig. 3.**
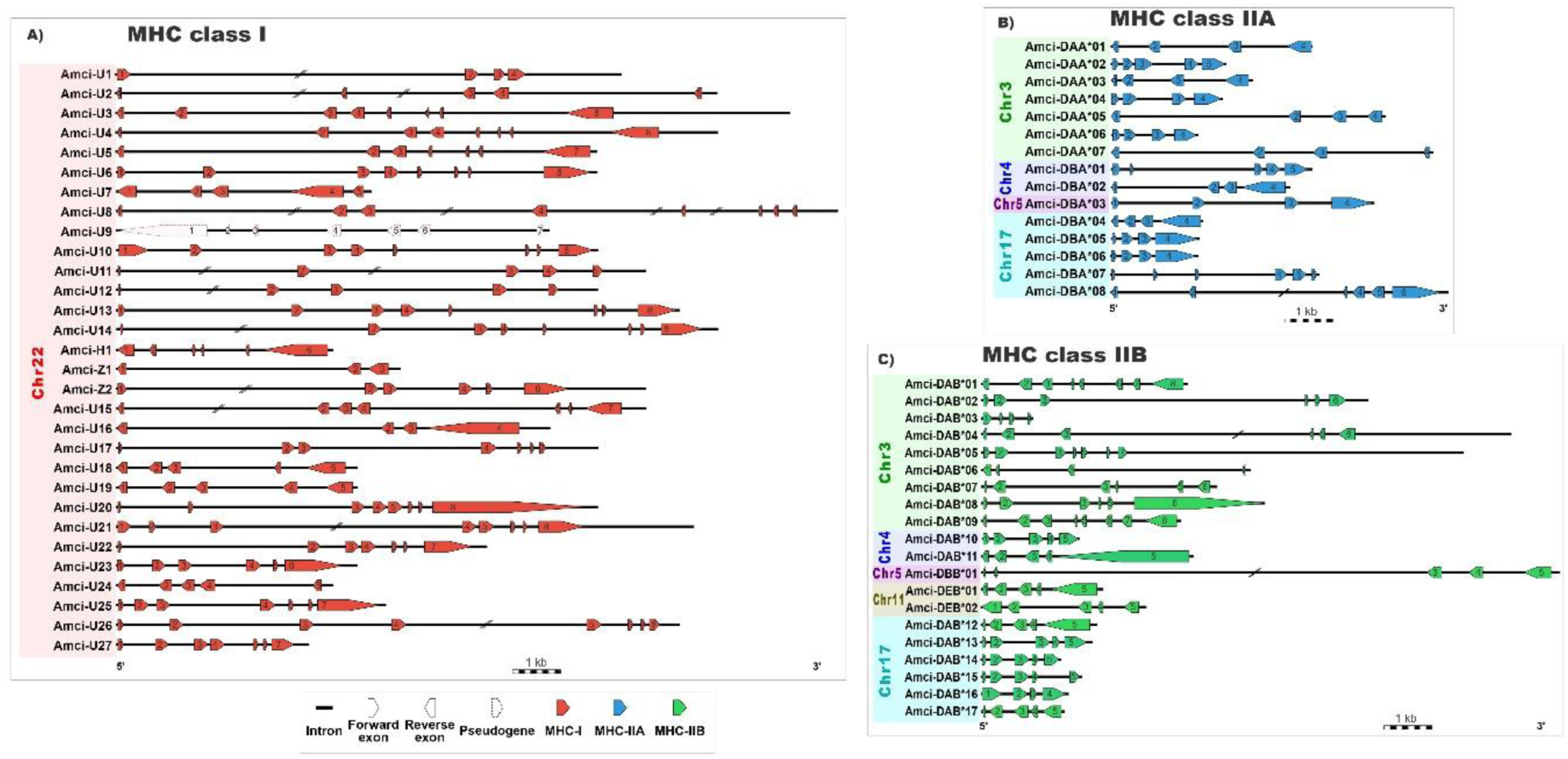
Structural organization of MHC genes in the Midas cichlid. Schematic representation of exon-intron organization in A) MHC-I, B) MHC-IIA, C) MHC-IIB genes. Horizontal lines represent introns, and colored arrows indicate exon orientation and relative length. Exon numbers are indicated within the arrows, dashed outlines identify pseudogenes, and diagonal breaks indicate large omitted intronic intervals. Colors distinguish gene classes.

Gene duplication patterns were classified into four categories based on sequence similarity and genomic location: singleton (genes without identified duplicates), tandem (adjacent duplicates), proximal (duplicates separated by a few intervening genes), and dispersed (duplicates located farther apart or on different chromosomes). Tandem duplication was the predominant category, followed by proximal and dispersed duplications, with no singletons identified (Fig. 4). MHC-I genes exhibited all three duplication modes, including dispersed duplications located within chromosome 22. In contrast, MHC-IIA and MHC-IIB genes were classified as tandem or proximal duplications across five chromosomes (Fig. 4).

**Fig. 4.**
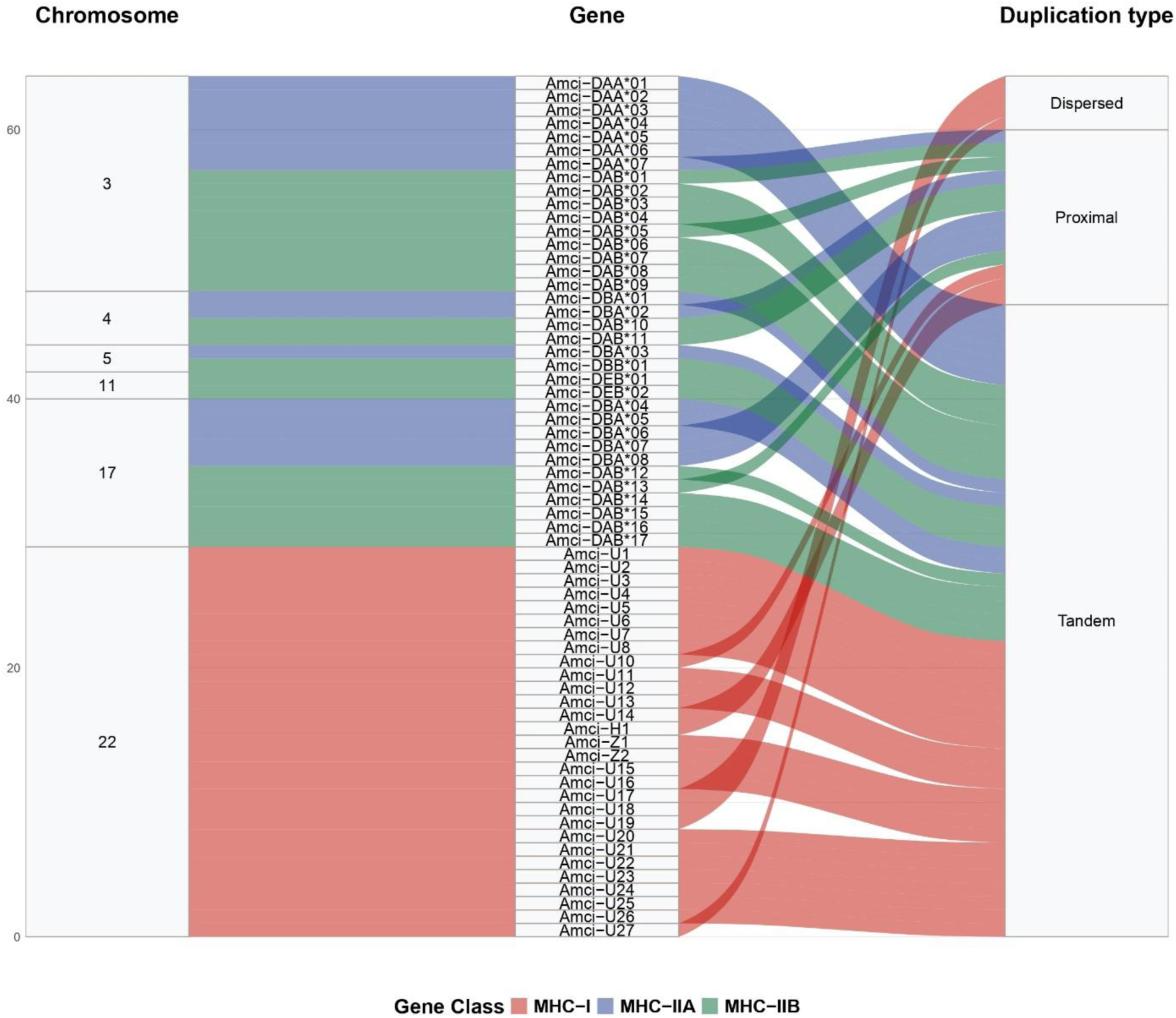
Duplication modes and chromosomal distribution of MHC genes in the Midas cichlid. Alluvial plot showing the relationships among chromosomal location, individual MHC genes, and duplication modes (tandem, proximal and dispersed). Flow widths are proportional to the number of genes, and colors distinguish MHC genes classes.

### Functional characterization and expression patterns of MHC genes

Functional domain analysis revealed that 83 of 156 MHC-related transcripts exhibited complete canonical domain architectures (Table S11). Among MHC-I transcripts, 13 of 45 showed partial or complete absence of key domains, including signal peptides, α1-α2 domains, Ig-like domains, transmembrane regions, and peptide-binding domains. Similar patterns were observed in seven of 18 MHC-IIA and seven of 25 MHC-IIB transcripts, primarily affecting signal peptides, class II α or β chain domains, transmembrane regions, and peptide-binding domains. In contrast, transcripts encoding antigen-processing components (*b2m*, *psmb5–9*, *tap1*, *tap2*, *tapbp*, and *tapbpl*), transcriptional regulators (*ciita* and *rfx*), and other associated genes (*brd2–4*, *brdt*, *cd74*, and *mHA*) generally retained their expected conserved domain architectures (Table S11).

Transcript expression varied across eight adult tissues (brain, eye, liver, lower pharyngeal jaw, muscle, skin, spleen, and thymus), with the highest observed expression levels in the immune-associated tissues, thymus and spleen (Fig. S5). Among MHC-I and MHC-II transcripts, Amci-DAA*06-P1, Amci-U3-P1, Amci-U6-P1, and Amci-DAB*02-P1 showed some of the highest observed expression levels, whereas other transcripts showed no detectable expression in the tissues examined. Among the remaining MHC-related genes, *b2m*, *psmb*, and *cd74* transcripts generally exhibited higher expression levels, while *mHA*, *tap*, *brd*, *ciita*, and *rfx* transcripts showed moderate to lower levels (Fig. S5).

### Transposable element distribution across MHC genomic regions

To characterize the TE distribution across MHC genomic regions, genome-wide TE annotations were intersected with four genomic compartments: 5-kb upstream and downstream flanking regions, exons, and introns. TEs were widely distributed across these compartments, although their coverage and composition varied among genomic regions (Fig. 5). Flanking regions and introns generally exhibited high TE coverage and broad representation of TE classes, while exonic regions showed a more restricted composition, predominantly comprising LINEs and DNA transposons. In comparison, antigen processing and regulatory genes generally exhibited lower TE coverage (Fig. S6).

**Fig. 5.**
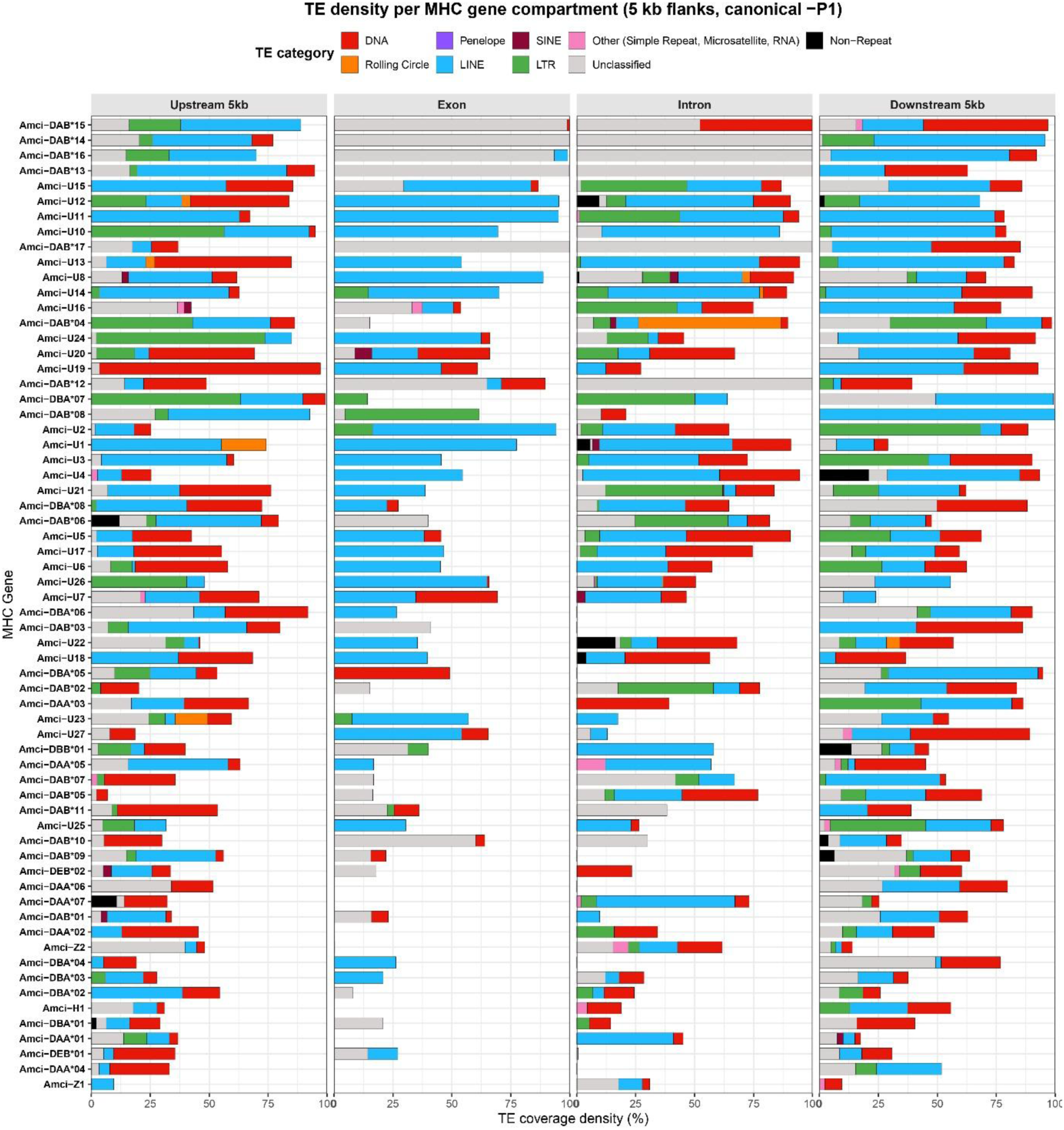
Transposable element (TE) distribution across MHC genes in the Midas cichlid. Stacked bar plots showing the coverage density of TE categories, expressed as a percentage of sequence length, across four genomic compartments: 5 kb upstream and downstream regulatory flanking regions, exons, and introns. Each row represents an MHC gene, with genes ordered by decreasing total TE coverage. Colors distinguish repetitive element categories.

Comparison with transcript expression profiles revealed descriptive differences among MHC-related members. Genes with relatively low TE coverage, including *b2m*, *psmb*, and *cd74*, generally showed higher expression levels across the tissues examined. In contrast, several MHC-I and MHC-II genes with greater TE coverage in flanking and intronic regions showed more variable expression across tissues, often with higher expression in the spleen and thymus. Some genes with relatively high exonic TE coverage also exhibited low transcript expression levels. Overall, these observations indicate that TE distribution and expression patterns differ among MHC-related genes, although a direct association between local TE coverage and gene expression remains to be established.

### Protein structural modeling

Candidate transcripts were selected for three-dimensional (3D) protein modeling based on complete functional domain architectures and relatively high expression levels, particularly in the spleen and thymus. The predicted MHC-I/*b2m* (Amci-U3-P1 + *b2m*-2-P1) and MHC-IIA/MHC-IIB (Amci-DAA*06-P1 + Amci-DAB*02-P1) complexes exhibited the expected MHC complex arrangements, with high overall model confidence (Fig. S7 and Table S13). The MHC-I/*b2m* complex had a mean predicted local distance difference test (pLDDT) score of 83.37, with 80.9% of residues scoring above 70, an interface predicted TM-score (ipTM) of 0.86, and a predicted TM score (pTM) of 0.78. The mean predicted aligned error (PAE) across extracellular domains was 11.9 Å, with higher values in transmembrane and terminal regions (Fig. S7a). The MHC-IIA/MHC-IIB complex had a higher mean pLDDT score of 88.32, with 87.9% of residues scoring above 70, an ipTM of 0.81, and a pTM of 0.80. Mean PAE across peptide-binding domains was 10.0 Å, with higher values in transmembrane and terminal regions (Fig. S7b).

### Phylogenetic relationships of MHC genes

ML phylogenies reconstructed from conserved peptide-binding domains revealed an association between phylogenetic clustering, chromosomal location, and gene duplication patterns (Fig. 6). For MHC-I, three major clades were identified, broadly reflecting the genomic organization of genes on chromosome 22, where tandem duplications predominated (Fig. 6a). The pseudogenes Amci-U9 and Amci-H1 were excluded because they lacked the conserved α1 and α2 domains required for phylogenetic analyses.

**Fig. 6.**
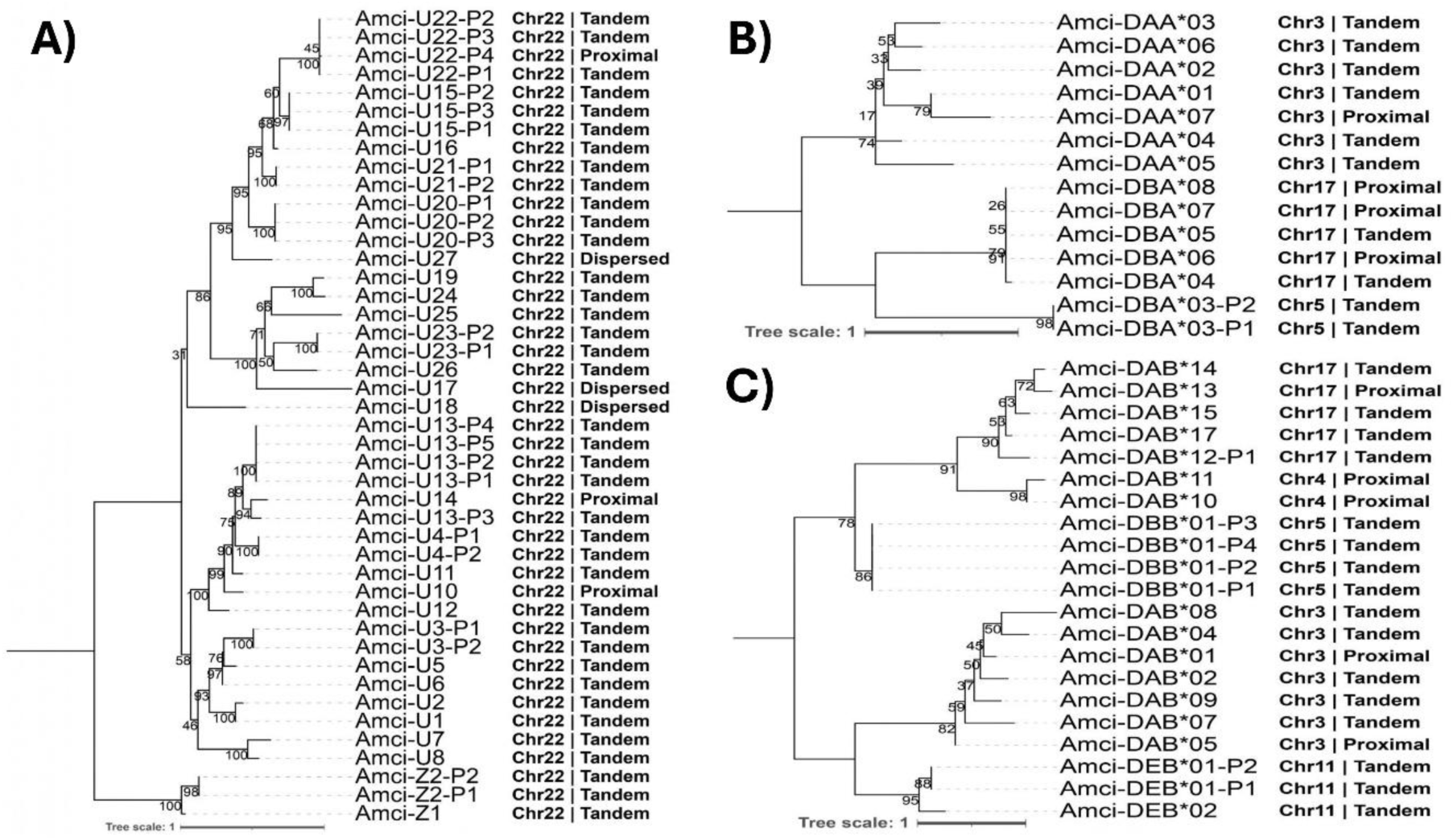
ML phylogenies of MHC class I and II genes in the Midas cichlid. Phylogenetic relationships were inferred from amino acid sequences of conserved domains: (A) MHC-I (α1-α2 domains; Pfam PF00129), (B) MHC-IIA (α1 domain; Pfam PF00993), and (C) MHC-IIB (β1 domain; Pfam PF00969 or Gene3D G3DSA:3.10.320.10). Tip labels indicate gene name, transcript variant, chromosomal location, and duplication mode (tandem, proximal, or dispersed). Bootstrap support values are shown at nodes. Trees are midpoint-rooted.

MHC-IIA sequences formed three major clades primarily associated with chromosomes 3, 5, and 17 (Fig. 6b). Gene located on chromosome 4 (Amci-DBA*01 and Amci-DBA*02) were excluded because no conserved α1 domain could be identified. The chromosome 5 clade comprised tandem duplicates, whereas clades associated with chromosome 3 and 17 included both tandem and proximal duplicates. MHC-IIB sequences formed four major clades associated with chromosomes 3, 5, 11, and a combined clade comprising genes from chromosomes 4 and 17 (Fig. 6c). Sequences lacking the conserved β1 domain (Amci-DAB*03, Amci-DAB*06, and Amci-DAB*16) were excluded. The chromosome 5 and 11 clades were predominantly associated with tandem duplications, whereas the chromosome 3 and 4/17 clades included both tandem and proximal duplicates. For sequences in the latter clade, the β1-domain was detected by Gene3D but not by Pfam.

An additional ML phylogeny combining previously published Midas cichlid MHC-IIB sequences (Hofmann et al. 2017; Bracamonte et al. 2022) revealed distinct phylogenetic groupings (Fig. S8). Sequences from chromosome 3 clustered with the DA lineage alongside all previously reported Midas MHC-IIB sequences. In contrast, sequences on chromosomes 4 and 17 formed a separate DA subclade, whereas those from chromosome 5 and 11 were assigned to the DB and DE lineages, respectively. This topology was consistent with the phylogenetic relationships inferred from the newly annotated sequences (Fig. 6 and S4).

### Analysis of positive selection and episodic diversification

To investigate patterns of selection across MHC genes, codon-based analyses were performed separately for peptide-binding (PBR) and non-PBR regions using site-level (SLAC, FEL, and MEME), gene-wide (BUSTED), and branch level (aBSREL) methods. BUSTED detected significant evidence of episodic positive selection in all three MHC classes and both functional partitions (p < 0.001), while aBSREL identified branches showing evidence of episodic positive selection across all classes (Table 1).

**Table 1.** Summary of positive selection analyses across PBR and non-PBR regions of MHC genes in the Midas cichlid. Codons indicate the number of aligned codon sites analyzed in each partition. FEL (Fixed Effect Likelihood) and MEME (Mixed Effects Model of Evolution) report the number of sites inferred to be under positive selection, with MEME results also expressed as percentage of the total codons analyzed. BUSTED reports p-values for gene-wide episodic positive selection, and aBSREL indicates the number of branches showing evidence of episodic positive selection relative to the total number of branches tested.

| MHC class | Region | Codons | FEL | MEME | BUSTED | aBSREL |
|---|---|---|---|---|---|---|
| MHC-I | PBR | 382 | 7 | 21 (5.50%) | 0.00 | 6/79 |
|  | non-PBR | 371 | 6 | 32 (8.63%) | 0.00 | 18/78 |
| MHC-IIA | PBR | 117 | 0 | 5 (4.27%) | 8.98 x 10<sup>-10</sup> | 1/23 |
|  | non-PBR | 206 | 1 | 8 (3.88%) | 5.22 x 10<sup>-6</sup> | 6/23 |
| MHC-IIB | PBR | 125 | 1 | 12 (9.60%) | 5.55 x 10<sup>-17</sup> | 6/37 |
|  | non-PBR | 283 | 3 | 11 (3.89%) | 4.79 x 10<sup>-5</sup> | 4/35 |

Site-level analysis using MEME, which detects episodic selection at individual codon sites, identified differences in the distribution of positively selected sites among MHC classes and functional partitions. In MHC-I, positive selection was detected at 21 PBR codons (5.50%) and 32 non-PBR codons (8.63%), with several sites showing high empirical Bayes posterior (EBP) support (≥ 0.90) and elevated estimates of the positive selection rate parameter (β+) (Fig. S9). In MHC-IIB, positively selected sites accounted for 9.60% of PBR codons, compared with 3.89% of non-PBR codons (Fig. S10). MHC-IIA showed lower proportions of positively selected sites in both partitions (4.27% in PBR and 3.88% in non-PBR; Fig. S11).

Overall, these analyses revealed contrasting patterns of molecular selection among MHC classes. MHC-I showed a higher partition of positively selected sites outside the PBR, while MHC-IIB exhibited a higher proportion with the PBR. MHC-IIA showed comparatively limited evidence of site-level positive selection, suggesting differences in selective pressures among MHC classes and functional regions.

## Discussion

This study provides a comprehensive, chromosome-resolved characterization of the MHC repertoire and associated antigen-processing-machinery in the Midas cichlid, representing one of the most detailed genomic analyses of the MHC currently available for cichlid fishes. By integrating genome annotation, transcriptomics, TE profiling, phylogenetic reconstruction, molecular selection analyses, and protein structural modeling, we revealed a highly expanded, multilocus, and chromosomally structured MHC repertoire characterized by extensive gene duplication and heterogeneous evolutionary pressures. These findings extend previous studies focused on individual MHC loci by providing a genome-wide perspective on MHC diversity, organization, and evolutionary dynamics, offering new insights into the genomic processes underlying immune repertoire diversification in cichlid fishes.

### Macroevolutionary dynamics and multilocus expansion of the MHC repertoire

At the genomic level, the Midas cichlid exhibits the dispersed organization of MHC-I and MHC-II genes characteristic of many teleost fishes (Bingulac-Popovic et al. 1997; Sato et al. 2000). MHC-I genes are concentrated on chromosome 22, whereas MHC-II genes are distributed across chromosomes 3, 4, 5, 11, and 17. This organization contrasts with the more tightly linked MHC regions characteristic of many tetrapods and cartilaginous fishes (Danchin et al. 2004; Belov et al. 2006; Shiina et al. 2017; Veríssimo et al. 2023; Hu et al. 2024). Chromosomal rearrangements and gene duplications, potentially associated with ancient vertebrate whole-genome duplication event and the teleost-specific duplication, have contributed to the diversification of immune gene organization across vertebrates (Sidow 1996; Meyer and Schartl 1999; Meyer and Van de Peer 2005). The genomic separation of MHC classes may also facilitate their evolutionary diversification under distinct selective and regulatory constraints, although these consequences remain to be directly evaluated in the Midas cichlid.

With 30 MHC-I, 15 MHC-IIA, and 20 MHC-IIB gene copies, the Midas cichlid exhibits a highly diversified repertoire with substantial representation of both MHC classes. A higher total number of MHC gene copies has been reported in another cichlid, the Nile tilapia (*Oreochromis niloticus;* 59 MHC-I, 33 MHC-IIA, and 16 MHC-IIB copies; Sato et al. 2006, 2012; Grimholt et al. 2015). These comparisons highlight the extensive MHC gene diversification observed in cichlids relative to several other surveyed teleosts, including sticklebacks (*Gasterosteus aculeatus;* 30 MHC-I, 11 MHC-IIA, and three MHC-IIB copies), zebrafish (*Danio rerio;* 30 MHC-I, 14 MHC-IIA, and seven MHC-IIB copies), Atlantic salmon (*Salmo salar;* 26 MHC-I, two MHC-IIA, and seven MHC-IIB copies), and medaka (*Oryzias latipes;* 18 MHC-I, six MHC-IIA, and five MHC-IIB copies) (Dijkstra et al. 2013; Grimholt et al. 2015, 2016). However, other teleost clades have followed markedly different evolutionary trajectories: Atlantic cod (*Gadus morhua;* Star et al. 2011), certain pipefishes (*Syngnathus* spp.; Haase et al. 2013; Small et al. 2016; Roth et al. 2020), and anglerfish (*Lophius piscatorius*; Dubin et al. 2019) have undergone secondary losses of the MHC-II pathway, sometimes accompanied by extensive MHC-I expansion, with up to 100 MHC-I copies reported in cod. Large MHC repertoires are not exclusive to teleosts, as some avian clades also exhibit extensive copy-number expansion, although often with marked asymmetry between MHC classes (Westerdahl et al. 2022). In contrast to teleost that have lost MHC-II, the Midas cichlid retains extensive repertoires of both MHC classes, highlighting the diversity of evolutionary trajectories underlying teleost antigen-presentation systems.

Phylogenetic analyses further reveal a heterogeneous representation of MHC-I and MHC-II lineages in the Midas cichlid genome. Most identified copies belonged to the classical lineages U (MHC-I) and DA (MHC-II), while additional genes were assigned to the non-classical lineages Z/H (MHC-I) and DB/DE (MHC-II). No representatives of the MHC-I L, S, or P lineages were detected (Dijkstra et al. 2007, 2013; Grimholt et al. 2015, 2019; Grimholt and Lukacs 2021). This uneven representation of MHC-I and MHC-II lineages, together with the observed patterns of gene expansion, is consistent with a birth-and-death model of multigene family evolution involving gene duplication, divergence, and loss (Nei and Rooney 2005). Nevertheless, the absence of detectable representatives of particular MHC-I lineages does not necessarily establish their evolutionary loss without broader comparative genomic evidence. Together, these findings suggest that duplication and differential retention of MHC genes have contributed to the heterogeneous composition of the Midas cichlid repertoire, although the evolutionary history of individual lineages remains to be resolved.

### Conservation of antigen-processing machinery and expression variation among MHC paralogs

The characterization of antigen processing and regulatory components (*b2m*, *tap*, *psmb*, *ciita*, *rfx*, and *cd74* members) revealed differences in genomic organization and expression patterns between MHC genes and their associated molecular machinery. In contrast to the extensive duplication and structural variation observed among MHC-I and MHC-II genes, antigen processing and regulatory genes generally exhibited more conserved gene architectures and fewer duplicated copies. Several of these genes also showed relatively broad expression across the tissues examined, particularly *bm2* and *cd74*. Together, these observations suggest that diversification of the MHC repertoire has occurred alongside the conservation of key components involved in antigen processing and regulation, potentially reflecting differences in functional constraints among gene families (Krawczyk et al. 2004; Seguín-Estévez et al. 2009; Yamaguchi and Dijkstra 2019).

Expression patterns also differed considerably among duplicated MHC genes. While some MHC-I and MHC-II paralogs exhibited relatively high expression in immune-associated tissues, particularly spleen and thymus, others showed low or undetectable expression across the tissues examined. Such expression differences are consistent with regulatory divergence following gene duplication and may reflect functional specialization, condition-dependent expression, or progressive loss of function (Lynch and Conery 2000; Innan and Kondrashov 2010; Sandve et al. 2018; Gerdol et al. 2019). The presence of truncated or partially disrupted MHC copies further suggests that potentially functional and degenerating genes coexist within the expanded repertoire, consistent with birth-and-death evolution. However, expression divergence alone does not establish subfunctionalization, and the functional status of individual paralogs will require further investigation.

The functional roles of different MHC-I and MHC-II lineages remain incompletely characterized in teleosts. In mammals, several non-classical MHC molecules perform specialized functions in immune regulation, while teleost MHC systems differ in aspects of antigen processing, including the general absence of the classical DM-mediated peptide-loading system (Alfonso and Karlsson 2000; Rodgers and Cook 2005; Dijkstra et al. 2013). In teleosts, the identification and classification of putatively non-classical MHC molecules therefore rely on phylogenetic relationships, sequence conservation, structural characteristics, and expression patterns, while their specific biological functions often remain experimentally unresolved (Grimholt 2016; Grimholt et al. 2019; Yamaguchi and Dijkstra 2019).

### Transposable element distribution and its potential implications for MHC repertoire evolution

Structural variation across the MHC repertoire is consistent with the dynamic evolution of this multigene family. Gene length, exon number, and intron size varied substantially, with MHC-IIB genes showing particularly pronounced structural variation. Variation in intron length may partly reflect differences in TE accumulation, while TE insertions and recombination-mediated rearrangements could also contribute to changes in gene architecture (Hofmann et al. 2017). In contrast, the relative conservation of exons encoding peptide-binding domains may reflect functional constraints associated with antigen presentation (Hughes and Nei 1989). Our analysis of TE distribution provides an additional perspective of the genomic context of MHC structural variation.

TEs were widely distributed across MHC genomic regions, with LINEs and DNA transposons contributing substantially to the repeat content of intronic and flanking regions. Such elements have been implicated in genomic rearrangements and gene-family diversification in cichlids (Carleton et al. 2020), although their direct contribution to MHC gene duplication in the Midas cichlid remains to be established. TE coverage and composition also varied among genomic compartments. Antigen-processing and regulatory genes generally exhibited lower TE coverage than MHC genes, suggesting that differences in TE distribution may reflect variation in genomic organization and functional constraints among MHC-related gene families. TE insertions within coding sequences can potentially disrupt gene function and contribute to pseudogenization (Bourque et al. 2018; Almeida et al. 2022). However, the contributions of these processes to MHC gene diversification and degeneration in the Midas cichlid remain unresolved and require further comparative functional investigation.

### Phylogenetic structure and genomic context of the MHC-IIB diversity

Phylogenetic analyses revealed a strong concordance between gene tree relationships and chromosomal location, suggesting that local duplication has contributed to the diversification of the MHC repertoire. For MHC-I, distinct phylogenetic clusters closely mirrored the spatial arrangement of genes along chromosome 22, while MHC-IIA and MHC-IIB genes exhibited chromosome-associated clustering patterns. These findings are consistent with the expansion and subsequent divergence of locally duplicated genes, although additional analyses would be required to determine whether gene conversion or concerted evolution has also contributed to their evolutionary history (Ohta 1991; Vinkler et al. 2023).

This genomic perspective is particularly relevant to previous population-level studies of MHC diversity in the Midas cichlid, which have primarily focused on sequence variation in exon 2 of MHC-IIB genes (Hofmann et al. 2017; Bracamonte et al. 2022). MHC-IIB genes located on chromosome 3 clustered within the DA lineage and were phylogenetically associated with the sequences reported in these earlier studies, while additional MHC-IIB copies were assigned to the DA, DB, and DE lineages. These findings demonstrate that the MHC-IIB sequences characterized in previous ecological and evolutionary studies represent only part of a broader, genomically distributed repertoire. Importantly, the presence of multiple related loci suggests that sequence diversity recovered using exon-based approaches may reflect a combination of allelic polymorphism and variation among paralogous genes, depending on the loci amplified and the resolution of the methods employed. Consequently, the chromosome-resolved genomic framework provided here offers an important reference for interpreting MHC-IIB diversity in natural populations and distinguishing allelic variation from differences in locus composition.

Phylogenetic relationships among MHC sequences from different species may be compatible with trans-species polymorphism (TSP), whereby allelic lineages persist across speciation events (Klein 1987; Klein et al. 2007; Lozano-Martín et al. 2023). However, similar phylogenetic patterns may also reflect the retention of ancient, duplicated loci rather than the persistence of ancestral allelic polymorphism. Distinguishing between these alternative explanations will require broader comparative analyses and a more detailed resolution of orthologous and paralogous relationships.

More broadly, the extensive MHC repertoire identified in the Midas cichlid extends genomic evidence of MHC expansion beyond the African cichlid species previously investigated (Málaga-Trillo et al. 1998; Figueroa et al. 2000; Sato et al. 2006, 2012; Blais et al. 2007; Hablützel et al. 2014). Together with the chromosome-level organization and phylogenetic diversity described here, these findings provide a foundation for comparative studies aimed at determining which features of MHC evolution are shared across cichlid lineages and which represent lineage-specific patterns of diversification.

### Domain-specific selection pressures and structural conservation

Analyses of molecular selection revealed marked differences in selective pressures among MHC gene families. In MHC-IIB, sites showing evidence of episodic positive selection were predominantly located within the PBR, consistent with the importance of this region in antigen recognition (Hughes and Nei 1989; Spurgin and Richardson 2010). In contrast, MHC-IIA exhibited comparatively limited evidence of positive selection, with 65% of the analyzed transcripts showing no detected positively selected sites. This asymmetry may reflect differences in functional constraints between MHC-II α and β chains, particularly their respective contributions to peptide binding and heterodimer formation (Brown et al. 1993; Dijkstra et al. 2013). The relatively conserved nature of MHC-IIA sequences may help maintain structural compatibility with divergent MHC-IIB partners, although the extent of heterodimer pairing flexibility in the Midas cichlid remains to be experimentally established. Differences in evolutionary constraints between MHC-IIA and MHC-IIB chains have also been discussed in birds (Westerdahl et al. 2022).

In MHC-I, positively selected sites were detected both within and outside the PBR, with a higher proportion identified in non-PBR (8.63%) than in the PBR (5.50%). This pattern suggests that selective pressures may extend beyond residues directly involved in peptide binding. Non-PBR regions include domains involved in interactions with other components of the immune system, such as *b2m* and the CD8 co-receptor (Bjorkman et al. 1987; Salter et al. 1989; Cole et al. 2012). Consequently, selection acting in these regions could potentially influence protein stability of molecular interactions, although the specific functional consequences of the detected substitutions remain unknown. Further analyses mapping positively selected sites onto individual structural domains would help clarify whether these signals are associated with particular interaction interfaces.

Finally, three-dimensional protein structure predictions suggest that substantial sequence divergence has occurred alongside the retention of broadly conserved MHC structural features. Despite variation in sequence composition and gene architecture, predicted MHC-I and MHC-II structures generally retained the characteristic folds associated with antigen-presenting molecules (Bjorkman et al. 1987). Structural models of MHC chains and associated components, including *b2m*, provide additional support for the conservation of key architectural features of antigen-presentation machinery. However, structural similarity and high model confidence do not establish protein functionality, peptide-binding capacity, or the stability of specific molecular interactions. Together, these results suggest that MHC sequence diversification in the Midas cichlid has occurred within a broadly conserved structural framework, while the functional implications of individual substitutions and gene copies remain to be determined.

## Conclusions

This study provides the first comprehensive genomic characterization of the MHC repertoire in a Neotropical cichlid fish. By integrating chromosome-level genome annotation, MHC-I and MHC-II lineage assignment, TE profiling, transcriptomics, phylogenetic reconstruction, structural modeling, and codon-based selection analyses, we revealed an extensive, chromosomally structured MHC repertoire in the Midas cichlid, characterized by extensive gene duplication, phylogenetic diversification, and heterogeneous selective pressures. The distribution of TEs across MHC genomic regions suggests a potential contribution to the evolution of MHC architecture, although their direct involvement in gene duplication and pseudogenization remains to be established. Molecular selection analyses revealed evidence of positive selection in MHC-I and MHC-IIB genes, while MHC-IIA showed more limited evidence of positive selection. Despite extensive diversification, many MHC proteins retain characteristic domain architectures, while transcriptomic analyses reveal heterogeneous expression patterns across tissues, consistent with the potential conservation of antigen presentation functions in at least part of the repertoire.

By revealing the multilocus and chromosomally structured nature of MHC-IIB diversity, our study provides a genomic context for interpreting sequence variation previously documented in natural Midas cichlid populations. These findings provide a foundation for investigating the evolutionary dynamics of genes during the Midas cichlid radiation and establish a valuable reference for comparative immunogenomic studies across cichlid fishes.

## Methods

### Functional annotation of the Midas cichlid genome

The Neotropical Midas cichlid complex (*Amphilophus* spp.) is represented by three publicly available scaffold-level genome assemblies (GCA_015108585.1, GCA_000751415.1, and the structurally annotated assembly described by Lv et al. 2022) and one high-quality chromosome-level assembly (GCA_013435755.1) with structural annotation (Kautt et al. 2020). Compared to scaffold-level assemblies, the chromosome-level assembly provides greater contiguity and facilitates the identification and chromosomal localization of closely related paralogs. Furthermore, alternative haplotigs were separated into a secondary contig set using MARVEL (Kautt et al. 2020), reducing the risk of misidentifying allelic variants as paralogous loci. We therefore selected this reference genome for comprehensive functional annotation and characterization of the MHC repertoire.

Assembly quality and contiguity were assessed with QUAST v5.3.0 (Gurevich et al. 2013). Structural and functional annotation was generated using the NCBI Eukaryotic Genome Annotation Pipeline (EGAPx) v0.4.1-alpha (Thibaud-Nissen et al. 2016), specifying NCBI TaxID 61818 (*Amphilophus*). EGAPx integrates transcriptomic evidence, protein homology information, and *ab initio* gene prediction using Gnomon to generate gene models. Transcriptomic evidence for gene prediction was obtained from publicly available short-read RNA-seq datasets in the NCBI Sequence Read Archive (SRA), covering multiple adult tissues (spleen, liver, thymus, muscle, intestine, gills, pharyngeal jaw, skin, brain, and eye), whole-body larvae, and several species within the Midas complex; both paired-end and single-end datasets were included (Table S2). EGAPx also incorporates repeat-masking and transcript-evidence filtering procedures intended to reduce spurious gene predictions, although annotation of highly similar paralogs and repeat-rich regions may remain challenging.

Annotation completeness and gene model quality were assessed using GAQET2 v1 (Garcia-Carpintero et al. 2025), integrating AGAT v1.4.1 (Dainat et al. 2024) for structural annotation statistics, BUSCO v5.8.3 (Actinopterygii_odb10 dataset; Manni et al. 2021) and OMArk v0.3.1 (LUCA.h5 database, Cichlidae; Nevers et al. 2024) for gene-space completeness; PSAURON v1.0.4 (Sommer et al. 2025) for protein-coding gene model quality; and TEsorter v1.4.7 (Zhang et al. 2022), combined with DeTEnGA, for identification and classification of putative transposable element (TE)-derived sequences in the predicted proteome.

Transposable elements in the reference assembly were identified, classified, and annotated *de novo* using Earl Grey v7.3.1 (Baril et al., 2024), which combines RepeatModeler2 (Flynn et al. 2020) for repeat discovery with RepeatMasker (Smit et al. 2013) and the RepBase and Dfam 4.0 databases for classification and annotation. The pipeline also uses consensus refinement, automated curation, and iterative BLAST-based searches, to detect full-length and fragmented TE insertions. Earl Grey was run with Teleostei as the reference taxonomic group, 16 threads, TE library clustering to reduce redundancy, and filtering of putative spurious TE annotations shorter than 100 bp.

### Identification and characterization of MHC loci

To identify the MHC-related gene repertoire in the Midas cichlid genome, two complementary approaches were used. First, amino acid sequences of 173 MHC-related genes were retrieved from the well-characterized human MHC system (GRCh38.p14; GCF_000001405.40; Supplementary file S1), including genes from the main MHC region on chromosome 6 (6p21.3; Shiina et al. 2017) and paralogous regions on chromosomes 1, 5, 9, 12, 14, 15, 16, 17, 19, and 20 (Veríssimo et al. 2023). These sequences were analyzed together with the EGAPx-derived Midas cichlid proteome using OrthoFinder v2.5.5 (Emms and Kelly 2019) with default parameters. Midas cichlid proteins belonging to orthogroups containing human MHC reference sequences were identified and cross-referenced with the corresponding GFF3 annotation.

Second, a custom amino acid database was compiled from published Midas cichlid MHC class IIB sequences (Hofmann et al. 2017; Bracamonte et al. 2022; Lozano-Martín et al. 2023), and annotated MHC protein sequences from additional Neotropical and African cichlid species. These were retrieved from RefSeq-annotated assemblies of *Archocentrus centrarchus* (GCF_007364275), *Astatotilapia calliptera* (GCF_900246225.1), *Haplochromis burtoni* (GCF_018398535.1), *Maylandia zebra* (GCF_000238955), *Neolamprologus brichardi* (GCF_000239395.1), *Oreochromis aureus* (GCF_013358895.1), *Oreochromis niloticus* (GCF_001858045), *Pelmatolapia mariae* (GCF_036321145.2), *Pundamilia nyererei* (GCF_000239375.1), and *Simochromis diagramma* (GCF_900408965.1), as well as the submitter-annotated assembly of *Sarotherodon galilaeus* (GCA_046562855.1). The resulting dataset of 1,783 sequences (Supplementary file S2) was used as queries in BLASTP and TBLASTN searches against the EGAPx-derived proteome and genome assembly, respectively, using BLAST+ (Camacho et al. 2009).

Candidate hits were manually inspected and mapped to the genome annotation and assembly using Geneious Prime 2025.2.2 (Kearse et al. 2012). BLAST hits were converted to GFF format and imported as custom annotation tracks. Hits matching annotated MHC-related genes were retained, while those matching genes with non-MHC annotations were excluded. Hits associated to uncharacterized gene models were further evaluated using online NCBI BLASTP and TBLASTN searches to confirm their putative MHC identity.

For each identified MHC-related locus, including MHC class I, IIA, IIB genes; antigen processing components (*b2m*, *psmb5*, *psmb6*, *psmb7*, *psmb8*, *psmb9*, *tap1*, *tap2*, *tapbp*, and *tapbpl*); transcriptional regulators (*ciita* and *rfx*); and associated genes (*brd2*, *brd3*, *brd4*, *brdt*, *cd74*, and *mHA*), genomic coordinates and gene models were extracted. Gene and coding sequence (CDS) lengths, exon and intron counts and lengths, and descriptive statistics, (minimum, maximum, mean (x̅), standard deviation (SD), and coefficient of variation (CV = (SD/x̅) x 100)) were calculated.

Gene duplication patterns among MHC class I, IIA, and IIB genes were investigated using pairwise protein similarities obtained with DIAMOND BLASTP v2.1.11 (Buchfink et al. 2021). Alignment scores and genomic coordinates were analyzed using MCScanX (Wang et al. 2012) to identify collinear blocks and classify genes as singleton (without detected duplicates), proximal duplicates (separated by maximum 10 intervening genes), tandem duplicates (adjacent genes), and dispersed duplicates (located elsewhere on the same or different chromosomes). Genomic organization and duplication patterns were visualized using an alluvial plot generated in R with ggplot2 (Wickham 2016) and ggalluvial (Brunson 2020), linking MHC class, duplication category, and chromosomal location.

### MHC lineage assignment and nomenclature

To identify the evolutionary lineages represented in the Midas cichlid MHC class I and class II repertoires, maximum likelihood (ML) phylogenies were reconstructed using reference sequences from major vertebrates lineages, including Chondrichthyes, early-diverging actinopterygian lineages, and teleosts representing Anguilliformes, Clupeiformes, Salmoniformes, and Cypriniformes. Sarcopterygian and tetrapod sequences were included as outgroups (Table S8). Reference sequences were selected from curated datasets with established lineage assignments (Bannai and Nonaka 2013; Dijkstra et al. 2013; Grimholt et al. 2015, 2019; Grimholt and Lukacs 2021).

Full-length amino acid sequences were aligned with MAFFT v7.505 (Katoh and Standley 2013) with the L-INS-I strategy, and trimmed using trimAl v1.4.1 (Capella-Gutiérrez et al. 2009) with the automated heuristic method. ML phylogenies were inferred using IQ-TREE 2 v2.1.3 (Minh et al. 2020), with substitution models selected automatically by ModelFinder (Kalyaanamoorthy et al. 2017). Branch support was assessed using 1,000 ultrafast bootstrap replicates (UFBoot2; Hoang et al. 2018) and SH-like approximate likelihood radio tests (SH-aLRT). Identical amino acid sequences were assigned unique numerical suffixes to preserve individual gene copies during phylogenetic inference. Trees were visualized and annotated using iTOL v6 (Letunic and Bork 2024).

Midas cichlid MHC genes were named following established nomenclature guidelines (Klein et al. 1990; Grimholt 2016) and the IPD-MHC conventions for teleost fishes. Class I gene names comprised the species’ abbreviation (*Amci*), lineage designation (U, Z, S, L, P, or H), and a number indicating the genomic order. Class II names included *Amci*, the letter D (Duo), lineage designation (A, B, D, or E), chain type (A for α and B for β), and an asterisk followed by a two digits indicating genomic order. Alternative transcripts from the same gene were distinguished using either “transcript variant ##” or the suffix “-P#”.

### Functional characterization, expression profiling, and structural modeling of MHC genes

Functional potential and protein domain architecture were assessed for all transcribed MHC-related loci. Amino acid sequences of MHC class I, IIA, and IIB proteins, antigen processing components (*b2m*, *psmb5–9*, *tap1*, *tap2*, *tapbp*, *tapbpl*); transcriptional regulators (*ciita*, *rfx*); and associated genes (*brd2–4*, *brdt*, *cd74*, *mHA*) were analyzed using the InterProScan v107.0 (Blum et al. 2025). Domain architecture was examined for characteristic features, including peptide-binding and immunoglobulin-like domains, signal peptides, and transmembrane regions.

Transcript expression was assessed using publicly available RNA-seq datasets from the NCBI SRA, including eight adult tissues: brain, eye, liver, lower pharyngeal jaw, muscle, skin, spleen, and thymus (Table S12). Reads were quality-checked using FastQC v0.12.1 (Andrews 2017), summarized with MultiQC v1.24.1 (Ewels et al. 2016), and trimmed using fastp v0.22.0 (Chen et al. 2018). Transcript abundance was quantified against the complete coding transcriptome using Salmon v1.10.2 (Patro et al. 2017). Estimated counts and transcripts per million (TPM) values were imported using tximport (Soneson et al. 2016). Mean TPM values were calculated per tissue and log_2_-transformed (log_2_(TPM+1)). Analyses and visualizations were performed in R using tidyverse (Wickham et al. 2019), ggrepel, and DESeq2 (Love et al. 2014).

Transposable element (TE) distribution across MHC loci was characterized using four genomic compartments: upstream (5 kb from the transcription start site), exons, introns, and downstream (5 kb from the transcription termination site). Overlaps between Earl Grey TE annotations and these compartments were identified using BEDTools intersect v2.31.0 (Quinlan and Hall 2010). Overlapping fragments were classified into nine categories: DNA transposons, rolling circle transposons (RC/Helitrons), Penelope-like elements, long interspersed nuclear elements (LINEs), short interspersed nuclear elements (SINEs), long terminal repeat (LTR) retrotransposons, other repetitive elements (simple repeats, low-complexity regions, microsatellites, and non-coding RNAs), unclassified TEs, and non-repetitive genomic regions. Overlapping intervals were merged to calculate non-redundant sequence coverage (kb) per compartment. Coverage values were normalized and visualized using tidyverse packages.

MHC class I, IIA, IIB, and *b2m* sequences were selected for three-dimensional (3D) modelling based on complete domain architecture and high expression in lymphoid tissues (spleen and thymus). *b2m* was included as the obligate light-chain subunit required for classical MHC class I heterodimer folding and surface presentation. Individual protein structures were predicted using AlphaFold2 (Jumper et al. 2021), and heterodimers using AlphaFold-Multimer, both implemented through ColabFold v1.5.5 (Mirdita et al. 2022) with default parameters. Predicted structures were visualized using PyMOL v3.1.6.1. All models were generated without bound peptide ligands and interpreted as predictions of overall protein fold and domain organization rather than peptide-bound conformations.

### Phylogenetic analysis of MHC genes

Evolutionary relationships among Midas cichlid MHC genes were reconstructed separately for each class using their peptide-binding domains, identified through InterProScan annotations. For MHC class I, the Pfam domain PF00129 (α1+α2) was used; for class IIA, PF00993 (α1); for class IIB, both Pfam (PF00969: β1) and Gene3D G3DSA:3.10.320.10, which also detected highly divergent β1 domains missed by Pfam. Domain sequences were extracted and analyzed using the alignment, trimming, and ML phylogenetic workflow described above for lineage assignment.

An additional MHC class IIB phylogeny was reconstructed to integrate newly identified MHC-IIB loci with previously published sequences (Hofmann et al. 2017; Bracamonte et al. 2022). The β1 domains of published 210-amino acid sequences were identified using InterProScan and extracted. These sequences were combined with the newly predicted β1-domains and analyzed using the same ML phylogenetic workflow.

### Analysis of selection pressures in MHC genes

To compare selection pressures between functional regions of MHC genes, CDS of MHC class I, IIA, and IIB genes were aligned in-frame using TranslatorX (Abascal et al. 2010), with MAFFT. Alignments were partitioned into peptide-binding (PBR) and non-PBR regions based on InterProScan domain annotations, preserving codon reading frames and masking premature stop codons as gaps (-). Selection was assessed using nonsynonymous-to-synonymous substitution rate ratios (d_N_/d_S_, ω) in HyPhy v2.5.10 (Pond et al. 2020) with previously inferred ML gene trees as phylogenetic inputs. Analyses included site-level methods (SLAC, FEL, MEME), the branch-level method aBSREL, and the gene-wide tests BUSTED. Results were summarized to compare selection patterns between PBR and non-PBR regions. For sites showing evidence of episodic positive selection with MEME (p ≤ 0.05), empirical Bayes posterior (EBP) probabilities were examined to identify branches potentially under positive selection. Site-branch combinations with EBP ≥ 0.50 were retained for further characterization. Selection patterns were visualized with ggplot2.

## Acknowledgements

This research project was supported by access to the FinisTerrae III supercomputer and associated data storage infrastructure provided by the Galician Supercomputing Center (CESGA), an infrastructure funded through the NextGeneration EU 2021 Recovery, Transformation and Resilience Plan (ICT2021-006904), the Pluriregional Operational Programme of Spain 2014-2020 of the European Regional Development Fund (ERDF; ICTS-2019-02-CESGA-3), and the State Programme for the Promotion of Scientific and Technical Research and Innovation 2013-2016 (ERDF, CESG15-DE-3114). We also acknowledge the access to the DRAGO supercomputer and related technical support, provided by Spanish National Research Council (CSIC). Finally, we thank two anonymous reviewers for their valuable comments and suggestions, which helped improve the manuscript.

## Data availability

The functional genome annotation data generated in this study have been deposited in Zenodo https://doi.org/10.5281/zenodo.23695154. This dataset includes the complete annotation files, predicted protein and transcript sequences, and the configuration parameters used for the annotation of the *Amphilophus citrinellus* reference genome (GCA_013435755.1).

## References

Abascal F, Zardoya R, Telford MJ. 2010. TranslatorX: multiple alignment of nucleotide sequences guided by amino acid translations. Nucleic Acids Res. 38:W7–W13 10.1093/nar/gkq291.

Alfonso C, Karlsson L. 2000. Nonclassical MHC class II molecules. Annu Rev Immunol. 18:113–142. 10.1146/annurev.immunol.18.1.113.

Almeida MV, Vernaz G, Putman ALK, Miska EA. 2022. Taming transposable elements in vertebrates: from epigenetic silencing to domestication. Trends Genet. 38:529–553. 10.1016/j.tig.2022.02.009.

Andrews S. 2017. FastQC: a quality control tool for high throughput sequence data. https://www.bioinformatics.babraham.ac.uk/projects/fastqc/.

Bannai HP, Nonaka M. 2013. Comprehensive analysis of medaka major histocompatibility complex (MHC) class II genes: Implications for evolution in teleosts. Immunogenetics. 65:883–895. 10.1007/s00251-013-0731-8.

Baril T, Galbraith J, Hayward A. 2024. Earl Grey: a fully automated user-friendly transposable element annotation and analysis pipeline. Mol Biol Evol. 41:msae068 10.1093/molbev/msae068.

Barluenga M, Stölting KN, Salzburger W, Muschick M, Meyer A. 2006. Sympatric speciation in Nicaraguan crater lake cichlid fish. Nature. 439:719–723. 10.1038/nature04325.

Belov K et al. 2006. Reconstructing an Ancestral Mammalian Immune Supercomplex from a Marsupial Major Histocompatibility Complex. PLoS Biol. 4:e46. 10.1371/journal.pbio.0040046.

Bingulac-Popovic J et al. 1997. Mapping of Mhc class I and class II regions to different linkage groups in the zebrafish, *Danio rerio*. Immunogenetics. 46:129–134. 10.1007/s002510050251.

Bjorkman PJ et al. 1987. The foreign antigen binding site and T cell recognition regions of class I histocompatibility antigens. Nature. 329:512–518. 10.1038/329512a0.

Blais J et al. 2007. MHC adaptive divergence between closely related and sympatric African cichlids. PLoS One. 2:e734. 10.1371/journal.pone.0000734.

Blum M et al. 2025. InterPro: the protein sequence classification resource in 2025. Nucleic Acids Res. 53:D444–D456. 10.1093/nar/gkae1082.

Bourque G et al. 2018. Ten things you should know about transposable elements. Genome Biol. 19:199. 10.1186/s13059-018-1577-z.

Bracamonte SE, Hofmann MJ, Lozano-Martín C, Eizaguirre C, Barluenga M. 2022. Divergent and non-parallel evolution of MHC IIB in the Neotropical Midas cichlid species complex. BMC Ecol Evol. 22:41. 10.1186/s12862-022-01997-9.

Brown JH et al. 1993. Three-dimensional structure of the human class II histocompatibility antigen HLA-DR1. Nature. 364:33–39. 10.1038/364033a0.

Brunson JC. 2020. ggalluvial: layered grammar for alluvial plots. J Open Source Softw. 5:2017. 10.21105/joss.02017.

Buchfink B, Reuter K, Drost HG. 2021. Sensitive protein alignments at tree-of-life scale using DIAMOND. Nat Methods 18:366–368. 10.1038/s41592-021-01101-x.

Camacho C et al. 2009. BLAST+: Architecture and applications. BMC Bioinform. 10:1–9. 10.1186/1471-2105-10-421.

Capella-Gutiérrez S, Silla-Martínez JM, Gabaldón T. 2009. trimAl: a tool for automated alignment trimming in large-scale phylogenetic analyses. Bioinformatics. 25:1972–1973. 10.1093/bioinformatics/btp348.

Carleton KL et al. 2020. Movement of transposable elements contributes to cichlid diversity. Mol Ecol. 29:4956–4969. 10.1111/mec.15685.

Chen S, Zhou Y, Chen Y, Gu J. 2018. fastp: an ultra-fast all-in-one FASTQ preprocessor. Bioinformatics. 34:i884–i890. 10.1093/bioinformatics/bty560.

Cole DK et al. 2012. The molecular determinants of CD8 co-receptor function. Immunology. 137:139–148. 10.1111/j.1365-2567.2012.03625.x.

Dainat J et al. 2024. AGAT: Another Gff Analysis Toolkit to handle annotations in any GTF/GFF format. Version v1.4.1. Zenodo. https://zenodo.org/records/13799920.

Danchin E et al. 2004. The major histocompatibility complex origin. Immunol Rev. 198:216–232. 10.1111/j.0105-2896.2004.00132.x.

Dijkstra JM et al. 2007. A third broad lineage of major histocompatibility complex (MHC) class I in teleost fish; MHC class II linkage and processed genes. Immunogenetics. 59:305–321. 10.1007/s00251-007-0198-6.

Dijkstra JM, Grimholt U, Leong J, Koop BF, Hashimoto K. 2013. Comprehensive analysis of MHC class II genes in teleost fish genomes reveals dispensability of the peptide-loading DM system in a large part of vertebrates. BMC Evol Biol. 13:260. 10.1186/1471-2148-13-260.

Dubin A, Jørgensen TE, Moum T, Johansen SD, Jakt LM. 2019. Complete loss of the MHC II pathway in an anglerfish, *Lophius piscatorius*. Biol Lett. 15:20190594. 10.1098/rsbl.2019.0594.

Eizaguirre C, Lenz TL, Kalbe M, Milinski M. 2012. Rapid and adaptive evolution of MHC genes under parasite selection in experimental vertebrate populations. Nat Commun. 3:621. 10.1038/ncomms1632.

Eizaguirre C, Lenz TL, Traulsen A, Milinski M. 2009. Speciation accelerated and stabilized by pleiotropic major histocompatibility complex immunogenes. Ecol Lett. 12:5–12. 10.1111/j.1461-0248.2008.01247.x.

Emms DM, Kelly S. 2019. OrthoFinder: phylogenetic orthology inference for comparative genomics. Genome Biol. 20:238. 10.1186/s13059-019-1832-y.

Ewels P, Magnusson M, Lundin S, Käller M. 2016. MultiQC: summarize analysis results for multiple tools and samples in a single report. Bioinformatics. 32:3047–3048. 10.1093/bioinformatics/btw354.

Figueroa F et al. 2000. Mhc class II B gene evolution in East African cichlid fishes. Immunogenetics. 51:556–575. 10.1007/s002510000181.

Flynn JM et al. 2020. RepeatModeler2 for automated genomic discovery of transposable element families. Proc Natl Acad Sci U.S.A. 117:9451–9457. 10.1073/pnas.1921046117.

Garcia-Carpintero V, de Martín I, García-Juan S, Bombarely A. 2025. Structural genome annotation QC with GAQET2 v2. Protocols.io. 10.17504/protocols.io.8epv5k5rnv1b/v1.

Geraghty DE, Daza R, Williams LM, Vu Q, Ishitani A. 2002. Genetics of the immune response: Identifying immune variation within the MHC and throughout the genome. Immunol Rev. 190:69–85. 10.1034/j.1600-065X.2002.19006.x.

Gerdol M et al. 2019. Molecular and structural characterization of MHC class II β genes reveals high diversity in the cold-adapted icefish *Chionodraco hamatus*. Sci Rep. 9:5523. 10.1038/s41598-019-42003-5.

Germain RN. 1994. MHC-dependent antigen processing and peptide presentation: providing ligands for T lymphocyte activation. Cell. 76:287–299. 10.1016/0092-8674(94)90336-0.

Grimholt U et al. 2015. A comprehensive analysis of teleost MHC class I sequences. BMC Evo. Biol. 15:32. 10.1186/s12862-015-0309-1.

Grimholt U. 2016. MHC and evolution in teleosts. Biology. 5:6. 10.3390/biology5010006.

Grimholt U, Lukacs M. 2021. Fate of MHCII in salmonids following 4WGD. Immunogenetics. 73:79–91. 10.1007/s00251-020-01190-6.

Grimholt U, Tsukamoto K, Hashimoto K, Dijkstra JM. 2019. Discovery of a novel MHC class I lineage in teleost fish which shows unprecedented levels of ectodomain deterioration while possessing an impressive cytoplasmic tail motif. Cells. 8:1056. 10.3390/cells8091056.

Gurevich A, Saveliev V, Vyahhi N, Tesler G. 2013. QUAST: quality assessment tool for genome assemblies. Bioinformatics. 29:1072–1075. 10.1093/bioinformatics/btt086.

Haase D et al. 2013. Absence of major histocompatibility complex class II mediated immunity in pipefish, *Syngnathus typhle*: evidence from deep transcriptome sequencing. Biol Lett. 9: 20130044. 10.1098/rsbl.2013.0044.

Hablützel PI et al. 2014. Intermediate number of major histocompatibility complex class IIB length variants relates to enlarged perivisceral fat deposits in the blunt-head cichlid *Tropheus moorii*. J Evol Biol. 27:2177–2190. 10.1111/jeb.12467.

Hablützel PI, Grégoir AF, Vanhove MPM, Volckaert FAM, Raeymaekers JAM. 2016. Weak link between dispersal and parasite community differentiation or immunogenetic divergence in two sympatric cichlid fishes. Mol Ecol. 25:5451–5466. 10.1111/mec.13833.

Hoang DT, Chernomor O, von Haeseler A, Minh BQ, Vinh LS. 2018. UFBoot2: improving the ultrafast bootstrap approximation. Mol Biol Evol. 35:518–522. 10.1093/molbev/msx281.

Hofmann MJ, Bracamonte SE, Eizaguirre C, Barluenga M. 2017. Molecular characterization of MHC class IIB genes of sympatric Neotropical cichlids. BMC Genet. 18:15. 10.1186/s12863-017-0474-x.

Hu J et al. 2024. A new chromosome-scale duck genome shows a major histocompatibility complex with several expanded multigene families. BMC Biol. 22:31. 10.1186/s12915-024-01817-0.

Hughes AL, Nei M. 1989. Nucleotide substitution at major histocompatibility complex class II loci: evidence for overdominant selection. Proc Natl Acad Sci U S A. 86:958–62. 10.1073/pnas.86.3.958.

Innan H, Kondrashov F. 2010. The evolution of gene duplications: classifying and distinguishing between models. Nat Rev Genet. 11:97–108. 10.1038/nrg2689.

Janeway CA, Travers P, Walport M, Shlomchik MJ. 2001. The major histocompatibility complex and its functions. In: Charles A Janeway J, Travers P, Walport M, Shlomchik MJ, editors. Immunobiology: The Immune System in Health and Disease 5th ed. Garland Science; p. 2–154. https://www.ncbi.nlm.nih.gov/books/NBK27156/.

Jumper J et al. 2021. Highly accurate protein structure prediction with AlphaFold. Nature. 596:583–589. 10.1038/s41586-021-03819-2.

Kalyaanamoorthy S, Minh BQ, Wong TKF, von Haeseler A, Jermiin LS. 2017. ModelFinder: fast model selection for accurate phylogenetic estimates. Nat Methods. 14:587–589. 10.1038/nmeth.4285.

Kasahara M. 1999. The chromosomal duplication model of the major histocompatibility complex. Immunol Rev. 167:17–32. 10.1111/j.1600-065X.1999.tb01379.x.

Katoh K, Standley DM. 2013. MAFFT Multiple Sequence Alignment Software Version 7: Improvements in Performance and Usability. Mol Biol Evol. 30:772–780. 10.1093/molbev/mst010.

Kautt AF et al. 2020. Contrasting signatures of genomic divergence during sympatric speciation. Nature. 588:106–111. 10.1038/s41586-020-2845-0.

Kearse M et al. 2012. Geneious Basic: An integrated and extendable desktop software platform for the organization and analysis of sequence data. Bioinformatics. 28:1647–1649. 10.1093/bioinformatics/bts199.

Klein J. 1987. Origin of major histocompatibility complex polymorphism: The trans-species hypothesis. Hum Immunol. 19:155–162. 10.1016/0198-8859(87)90066-8.

Klein J et al. 1990. Nomenclature for the major histocompatibility complexes of different species: a proposal. Immunogenetics. 31:217–219.

Klein J, Sato A, Nikolaidis N. 2007. MHC, TSP, and the origin of species: From immunogenetics to evolutionary genetics. Annu Rev Genet. 41:281–304. 10.1146/annurev.genet.41.110306.130137.

Krawczyk M et al. 2004. Long distance control of MHC class II expression by multiple distal enhancers regulated by regulatory factor X complex and CIITA. J Immunol. 173:6200–6210. 10.4049/jimmunol.173.10.6200.

Letunic I, Bork P. 2024. Interactive Tree of Life (iTOL) v6: recent updates to the phylogenetic tree display and annotation tool. Nucleic Acids Res. 52:W78–W82. 10.1093/nar/gkae268.

Love MI, Huber W, Anders S. 2014. Moderated estimation of fold change and dispersion for RNA-seq data with DESeq2. Genome Biol. 15:550. 10.1186/s13059-014-0550-8.

Lozano-Martín C, Bracamonte SE, Barluenga M. 2023. Evolution of MHC IIB diversity across cichlid fish radiations. Genome Biol Evol. 15:evad110. 10.1093/gbe/evad110.

Lv Y et al. 2022. An updated genome assembly improves understanding of the transcriptional regulation of coloration in Midas cichlid. Front Mar Sci. 9:950573. 10.3389/fmars.2022.950573.

Lynch M, Conery JS. 2000. The Evolutionary Fate and Consequences of Duplicate Genes. Science. 290:1151–1155. 10.1126/science.290.5494.1151.

Málaga-Trillo E et al. 1998. Linkage relationships and haplotype polymorphism among cichlid Mhc class II B Loci. Genetics. 149:1527–1537. 10.1093/genetics/149.3.1527.

Malmstrøm M et al. 2016. Evolution of the immune system influences speciation rates in teleost fishes. Nat Genet. 48:1204–1210. 10.1038/ng.3645.

Manni M, Berkeley MR, Seppey M, Simão FA, Zdobnov EM. 2021. BUSCO update: novel and streamlined workflows along with broader and deeper phylogenetic coverage for scoring of eukaryotic, prokaryotic, and viral genomes. Mol Biol Evol. 38:4647–4654. 10.1093/molbev/msab199.

Meyer A, Schartl M. 1999. Gene and genome duplications in vertebrates: the one-to-four (-to-eight in fish) rule and the evolution of novel gene functions. Curr Opin Cell Biol. 11:699–704. 10.1016/S0955-0674(99)00039-3.

Meyer A, Van de Peer Y. 2005. From 2R to 3R: evidence for a fish-specific genome duplication (FSGD). BioEssays. 27:937–945. 10.1002/bies.20293.

Michalová V, Murray BW, Sültmann H, Klein J. 2000. A contig map of the Mhc class I genomic region in the zebrafish reveals ancient synteny. J Immunol. 164:5296–5305. 10.4049/jimmunol.164.10.5296.

Minh BQ et al. 2020. IQ-TREE 2: New models and efficient methods for phylogenetic inference in the genomic era. Mol Biol Evol. 37:1530–1534. 10.1093/molbev/msaa015.

Mirdita M et al. 2022. ColabFold: making protein folding accessible to all. Nat Methods 19:679–682. 10.1038/s41592-022-01488-1.

Murphy K, Weaver C. 2016. Janeway’s Immunobiology. 9th ed. Murphy K, Weaver C, editors. Garland Sciences, Taylor & Francis Group, LLC.

Neefjes J, Jongsma MLM, Paul P, Bakke O. 2011. Towards a systems understanding of MHC class I and MHC class II antigen presentation. Nat Rev Immunol. 11:823–836. 10.1038/nri3084.

Nei M, Rooney AP. 2005. Concerted and birth-and-death evolution of multigene families. Annu Rev Genet. 39:121–152. 10.1146/annurev.genet.39.073003.112240.

Nevers Y et al. 2024. Quality assessment of gene repertoire annotations with OMArk. Nat Biotechnol. 43:124–133. 10.1038/s41587-024-02147-w.

Ono H, O’hUigin C, Vincek V, Klein J. 1993. Exon-intron organization of fish major histocompatibility complex class II B genes. Immunogenetics. 38:223–234. 10.1007/BF00211522.

Ohta T. 1991. Multigene families and the evolution of complexity. J Mol Evol. 33:34–41. 10.1007/bf02100193.

Patro R, Duggal G, Love MI, Irizarry RA, Kingsford C. 2017. Salmon provides fast and bias-aware quantification of transcript expression. Nat Methods 14:417–419. 10.1038/nmeth.4197.

Pinesha N, Harmand TJ, Ploegh HL. 2022. A guide to antigen processing and presentation. Nat. Rev Immunol. 22:751–764. 10.1038/s41577-022-00707-2.

Pond SLK et al. 2020. HyPhy 2.5—a customizable platform for evolutionary hypothesis testing using phylogenies. Mol Biol Evol. 37:295–299, 10.1093/molbev/msz197.

Quinlan AR, Hall IM. 2010. BEDTools: a flexible suite of utilities for comparing genomic features. Bioinformatics. 26:841–842. 10.1093/bioinformatics/btq033.

Rodgers JR, Cook RG. 2005. MHC class Ib molecules bridge innate and acquired immunity. Nat Rev Immunol. 5:459–471. 10.1038/nri1635.

Ronco F et al. 2021. Drivers and dynamics of a massive adaptive radiation in cichlid fishes. Nature. 589:76–81. 10.1038/s41586-020-2930-4.

Roth O et al. 2020. Evolution of male pregnancy associated with remodeling of canonical vertebrate immunity in seahorses and pipefishes. Proc Natl Acad Sci U.S.A. 117:9431–9439. 10.1073/pnas.1916251117.

Salter RD et al. 1989. Polymorphism in the α3 domain of HLA-A molecules affects binding to CD8. Nature. 338:345–347. 10.1038/338345a0.

Salzburger W. 2018. Understanding explosive diversification through cichlid fish genomics. Nat Rev Genet. 19:705–717. 10.1038/s41576-018-0043-9.

Sandve SR, Rohlfs RV, Hvidsten TR. 2018. Subfunctionalization versus neofunctionalization after whole-genome duplication. Nat Genet. 50:908–909. 10.1038/s41588-018-0162-4.

Sato A et al. 2000. Nonlinkage of major histocompatibility complex class I and class II loci in bony fishes. Immunogenetics. 51:108–116. 10.1007/s002510050019.

Sato A et al. 2006. Mhc class I genes of the cichlid fish *Oreochromis niloticus*. Immunogenetics. 58:917–928. 10.1007/s00251-006-0151-0.

Sato A, Dongak R, Hao L, Shintani S, Sato T. 2012. Organization of Mhc class II A and B genes in the tilapiine fish *Oreochromis*. Immunogenetics. 64:679–690. 10.1007/s00251-012-0618-0.

Seguín-Estévez Q et al. 2009. The transcription factor RFX protects MHC class II genes against epigenetic silencing by DNA Methylation. J Immunol. 183:2545–2553. 10.4049/jimmunol.0900376.

Shiina T, Blancher A, Inoko H, Kulski JK. 2017. Comparative genomics of the human, macaque and mouse major histocompatibility complex. Immunology. 150:127–138. 10.1111/imm.12624.

Sidow A. 1996. Gen(om)e duplications in the evolution of early vertebrates. Curr Opin Genet Dev. 6:715–722. 10.1016/S0959-437X(96)80026-8.

Small CM et al. 2016. The genome of the Gulf pipefish enables understanding of evolutionary innovations. Genome Biol. 17:258. 10.1186/s13059-016-1126-6.

Smit AFA, Hubley R, Green P. RepeatMasker Open-4.0. 2013–2015. http://www.repeatmasker.org. Accessed 12 September 2026.

Sommer MJ, Zimin A V., Salzberg SL. 2025. PSAURON: a tool for assessing protein annotation across a broad range of species. NAR Genom Bioinform. 7:lqae189. 10.1093/nargab/lqae189.

Soneson C, Love MI, Robinson MD. 2016. Differential analyses for RNA-seq: transcript-level estimates improve gene-level inferences. F1000Res. 4:1521. 10.12688/f1000research.7563.2.

Spurgin LG, Richardson DS. 2010. How pathogens drive genetic diversity: MHC, mechanisms and misunderstandings. Proc R Soc Lond B Biol Sci. 277:979–988. 10.1098/rspb.2009.2084.

Star B et al. 2011. The genome sequence of Atlantic cod reveals a unique immune system. Nature. 477:207–210. 10.1038/nature10342.

Thibaud-Nissen F et al. 2016. P8008 The NCBI Eukaryotic Genome Annotation Pipeline. J Anim Sci. 94:184. 10.2527/jas2016.94supplement4184x.

Veríssimo A et al. 2023. An ancestral major histocompatibility complex organization in cartilaginous fish: reconstructing MHC origin and evolution. Mol Biol Evol. 40:msad262. 10.1093/molbev/msad262.

Vinkler M et al. 2023. Understanding the evolution of immune genes in jawed vertebrates. J Evol Biol. 36:847–873. 10.1111/jeb.14181.

Wang Y et al. 2012. MCScanX: a toolkit for detection and evolutionary analysis of gene synteny and collinearity. Nucleic Acids Res. 40:e49. 10.1093/nar/gkr1293.

Westerdahl H et al. 2022. The genomic architecture of the passerine MHC region: High repeat content and contrasting evolutionary histories of single copy and tandemly duplicated MHC genes. Mol Ecol Resour. 22:2379–2395. 10.1111/1755-0998.13614.

Wickham H. 2016. ggplot2: elegant graphics for data analysis. Springer-Verlag New York. 10.1007/978-3-319-24277-4.

Wickham, et al. 2019. Welcome to the Tidyverse. J Open Source Soft. 4:1686. 10.21105/joss.01686.

Wieczorek M et al. 2017. Major histocompatibility complex (MHC) class I and MHC class II proteins: Conformational plasticity in antigen presentation. Front Immunol. 8:292. 10.3389/fimmu.2017.00292.

Yamaguchi T, Dijkstra JM. 2019. Major Histocompatibility Complex (MHC) genes and disease resistance in fish. Cells. 8:378. 10.3390/cells8040378.

Zhang RG et al. 2022. TEsorter: An accurate and fast method to classify LTR-retrotransposons in plant genomes. Hortic Res. 9:uhac017. 10.1093/hr/uhac017.

Zinkernagel RM, Doherty PC. 1997. The discovery of MHC restriction. Immunol Today. 18:14–17. 10.1016/S0167-5699(97)80008-4.

